# Evaluating Antioxidant Capacity of Different Propolis Samples from Konya, Turkey and Their Inhibitory Effect on Head and Neck Cancer Cells

**DOI:** 10.1101/183913

**Authors:** Sultan Ciftci-Yilmaz, Zeynep N. Azman, Kubra Kosem, Esra Gunduz, Reidar Grenman

**Author notes:** Corresponding author: Sultan Ciftci-Yilmaz, Ph.D., Department of Genetic Research, Institute for Research and Medical Consultations, University of Dammam (Imam Abdulrahman Bin Faisal University), Dammam 31441, Kingdom of Saudi Arabia.

## Abstract

Propolis is a resinous mixture collected and used by the honey bees to construct and repair their hives. The component of propolis varies depend on the type of the plants collected. Propolis and its constituents have been subjected to many studies and are known for their antioxidant, antimicrobial and anticarcinogenic properties. In our study, antioxidant and antitumor capacity of propolis from Konya Sakyatan and Kiziloren regions were investigated. According to our result, Kızıloren propolis sample possesses higher antioxidant component and antioxidant capacity than Sakyatan sample. Accordingly, Kiziloren sample showed antiproliferative effect at much lower doses compared to the Sakyatan sample. Both samples effectively inhibited the migration of cancer cells at their determined IC_50_ dosages. Obtained data indicates that constituents of propolis can greatly vary from one sample to another even in the same region and propolis selections for cancer prevention and treatment studies should be carefully considered.

## 1. Introduction

Propolis, a resinous substance constitutes bee enzyme, wax and resin collected from plants, is produced by bees for constructing and repairing their hives. Since the ancient times, propolis has been used as a remedy for many medical conditions because of its antiseptic and local anesthetic properties (Sforcin 2016). In the last decades, propolis has been extensively studied for understanding its chemical composition and biological properties (Sforcin 2016; DAntas Silva et al. 2017; Krol et al. 2013; Silva-Carvalho et al. 2015). Today it is known that propolis possesses many different biological properties including antioxidant, antiviral, anti-parasitic, anti-inflammatory and antitumor activities (Sforcin 2016; DAntas Silva et al. 2017; Krol et al. 2013; Silva-Carvalho et al. 2015; Oses et al. 2016).

Many researchers demonstrated propolis’s ability to inhibit cancer progression by inducing apoptosis and cell-cycle arrest and reducing metastasis and invasion in both in vitro and in vivo studies (Watanebe et al. 2011; Chan et al. 2013; Frión-Herrera et al. 2015; Patel et al. 2016). Not exerting any major side effect in both mice and human subjects propolis administrated makes propolis a valuable potential anticancer agent (Sforcin et al. 1995; Sforcin et al. 2002; Jasprica et al. 2007; Henatsch et al. 2016). One main problem with use of propolis is lack of its standardization since propolis’s constituents and dependently its biological activities can greatly vary based on the region and its extraction method (Dantas Silva et al. 2017; Bankova 2005; Marcucci 1995).

The aim of this study was to evaluate antioxidant and antitumor properties of two different propolis samples collected from two areas separated only by 70 km at Konya, Turkey region.

## 2. Results and discussion

### 2.1. Antioxidant properties

Phenolic compounds are one of the most important phytochemicals and are linked with health benefits such as anti-cancer, anti-oxidant, anti-microbial and anti-inflammatory. Total phenolic and flavonoid contents of the propolis extracts were calculated as gallic acid and rutin equivalents, respectively (Table 1). KK propolis contained higher phenolic content (94.54 mgGAEs/g extract) and flavonoid (81.73 mgREs/g extract) compared to KS propolis (40.83 mgGAEs/g extract and 52.26 mgREs/g extract). Apparently, KK propolis had 2.3 times more total phenolics than KS propolis. Similarly, polyphenols were identified as major components in the propolis from different origins (Righi et al. 2013; Miguel et al. 2014). Again, the level of polyphenols in the propolis samples show variations depending on their geographic origin, botanical sources and the collection seasons. Similar approaches were indicated by several researches (Silva et al. 2012; Socha et al. 2015).

**Table 1.** Antioxidant properties of two propolis (KK propolis and KS propolis)^a^

| Parameters | KS Propolis | KK propolis |
| --- | --- | --- |
| Total phenolic content (mgGAEs/g extract) | 40.83±0.25 | 94.54±1.12 |
| Total flavonoid content (mgREs/g extract) | 52.26±0.18 | 81.73±0.17 |
| DPPH radical Scavenging Activity (mgTEs/g extract) | 116.71±3.15 | 306.23±0.55 |
| Ferric reducing power (mgTEs/g extract) | 185.25±1.03 | 461.52±5.60 |
| Cupric reducing power (mgTEs/g extract) | 187.55±0.55 | 448.73±0.88 |
| Phosphomolybdenum assay (mmolTEs/g extract) | 2.21±0.11 | 2.40±0.15 |
| Ferrous ion chelating activity (mgEDTAEs/g extract) | 10.12±0.32 | 10.40±0.52 |

KK, Konya Kizilören; KS, Konya Sakyatan; GAEs, gallic acid equivalents; REs, rutin equivalents; TEs, trolox equivalents; EDTAEs, EDTA equivalents.
<sup>a</sup>Values expressed are means ± SD of three parallel measurements.

DPPH (purple) is a stable free radical and is a widely used method of evaluating of radical scavenging activities. In DPPH assay, this radical is reduced by reacting with an antioxidant to yellow-colored diphenylpicrylhydrazine. The changes were spectrophometrically measured at 515-517 nm. Table 1 shows the scavenging effect on DPPH radicals of propolis samples. In accordance with the total phenolic content, KK propolis (306.23 mgTEs/g extract) exhibited higher radical scavenging activity than KS propolis (116.71 mgTEs/g extract). The results obtained in the current study are in accordance with many authors, who confirmed by strong correlation between the amount of phenolic and free radical scavenging activity of propolis samples (Ahn et al. 2009; Kalogeropoulos et al. 2009).

Reducing power assays are often used as an indicator of electron-donating activity, which is an important mechanism of antioxidant compounds. Therefore, FRAP and CUPRAC methods were applied to evaluate reducing power potentials of propolis samples and the results are depicted in Table 1. KK propolis had the strongest reductive ability in both FRAP (461.52 mgTEs/g extract) and CUPRAC assays (448.73 mgTEs/g extract). These results showed KK propolis to be almost three times more potent than KS propolis in these assays. Reduction potentials of propolis from different origins were reported in the literature (Aliyazicioglu et al. 2013; Dong et al. 2013; Miguel et al. 2014). Similar to FRAP and CUPRAC assays, phosphomolybdenum assay based on based on the reduction of Mo (VI) to Mo (V) by the antioxidant compounds and the formation of green phosphate/Mo (V) complex. From Table 1, the phosphomolybdenum activity was obtained with KK propolis (2.40 mmolTEs/g extract) very closely by KS propolis (2.21 mmolTEs/g extract).

Transition metal ions, such as iron and copper, are known as important pro-oxidant in lipid peroxidation thereby the metal ions can accelerate the generation of the new free radicals in this process. In this direction, chelating agents or chelators may be inhibit radical formations by stabilizing the metal ions. The chelating capacities of propolis samples were estimated by ferrozine assay. Similar to phosphosmolybdenum assay, the propolis samples exhibited similar chelating activities (10.40 mgEDTAEs/g extract for KK propolis and 10.12 mgEDTAEs/g extract for KS propolis). According to these results, the observed chelating activity may be explained by the presence non-phenolic chelators such as polypeptides and ascorbic acid in the propolis samples. Similar observations were made in the different propolis samples (Miguel et al. 2014; Geckil et al. 2005).

### 2.2. Phenolic components

Table 2 lists the phenolic compounds of propolis samples that were analyzed qualitatively and quantitatively by HPLC-DAD. Sixteen standard compounds were analyzed and 14 of them were detected in the tested extracts. Gallic and chlorogenic acid were not found in these samples. This is in agreement with the reported by Falcao et al. (2010) and Popova et al., (2004) that found the lower level of gallic acid in propolis from temperate zones. The results revealed that quercetin (7.41 and 32.7 mg/g extract), hesperidin (5.62 and 22.4 mg/g extract) and apigenin (3.32 and 10.83 mg/g extract) were the main phenolic constituents in both KK and KS propolis extracts, with different abundances. Generally, all phenolic components in KK propolis were higher than KS propolis. In this point, our data suggest that propolis from different regions have different levels of phenolic components. Biological activities of polyphenol-rich extracts from propolis show variations depending on its chemical composition. Similar results were observed for different origins of same countries including Portugal (Silva et al. 2012; Moreira et al. 2008; Miguel et al. 2010), Turkey (Aliyazıcıoglu et al. 2013), Algeria (Benhanifia et al. 2013), China (Dong et al. 2013), Poland (Socha et al. 2015) and Brazilia (Righi et al. 2013). The major components, namely quercetin, hesperidin and apigenin, have considerable biological activities such as antioxidant, anticancer, antibacterial (Wilmsen et al. 2005; Kumar, Pandey 2005; Liu et al. 2013). In fact, the propolis samples could be considered as a source of valuable components. The observed high antioxidant properties for KK propolis may be explained with the higher levels of these major constituents as compared to KS propolis.

**Table 2.**
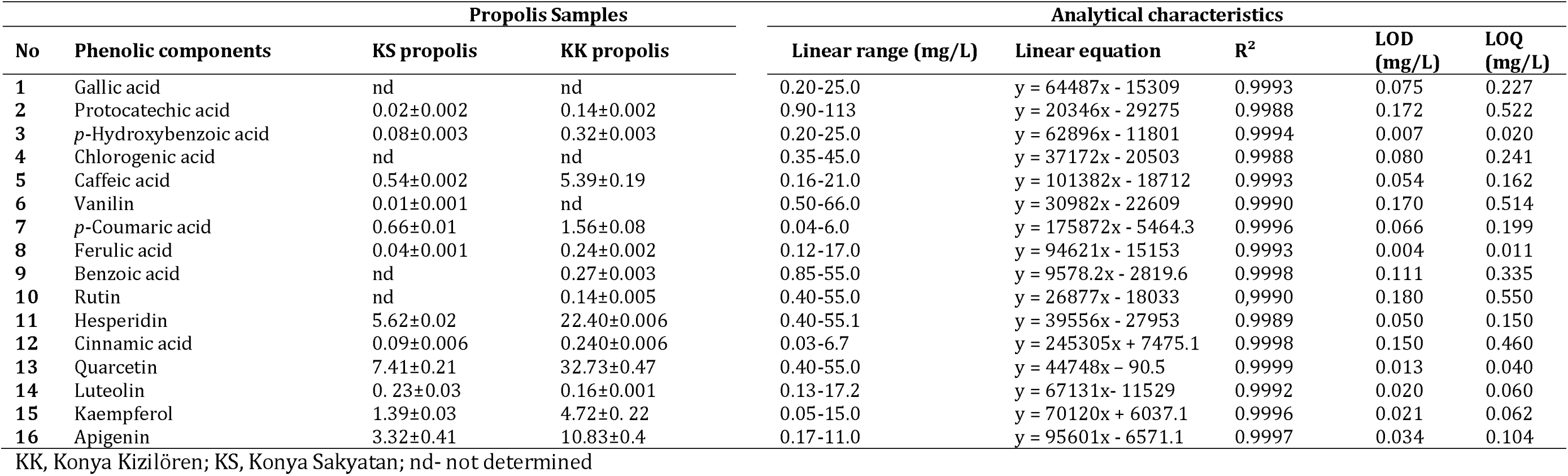
Phenolic components (mg/g extract) of two Propolis (mean ± SD).

### 2.3. Anticancer Properties

To assess the anticancer properties of KK and KS propolis samples, we used the UT-SCC-74A primary head and neck cancer cell line. Effect of both propolis on cell proliferation determined by measuring difference in cell index at six different propolis dosages (1000, 500, 250, 125, 31 and 0 μg/ml) by using xCELLigence real time cell analyzer system. First the cells were seeded to the E-plates. After allowing cells to attach for 24 hours, the cells were treated with different concentrations of propolis. Cell index was measured at every 15 minutes for 60 hours. Data analysis was performed using RTCA Software version 1.2. Based on the analysis, IC_50_ of KK was calculated as 39,12 μg/ml and IC_50_ of KS sample was calculated as 119,36 μg/ml at 24 hour following the propolis administration (Figure 1). The difference between the IC_50_ dosages of KK and KS propolis was correlated with their antioxidant properties and amount of the phenolic components (Table 1-2).

**Figure 1.**
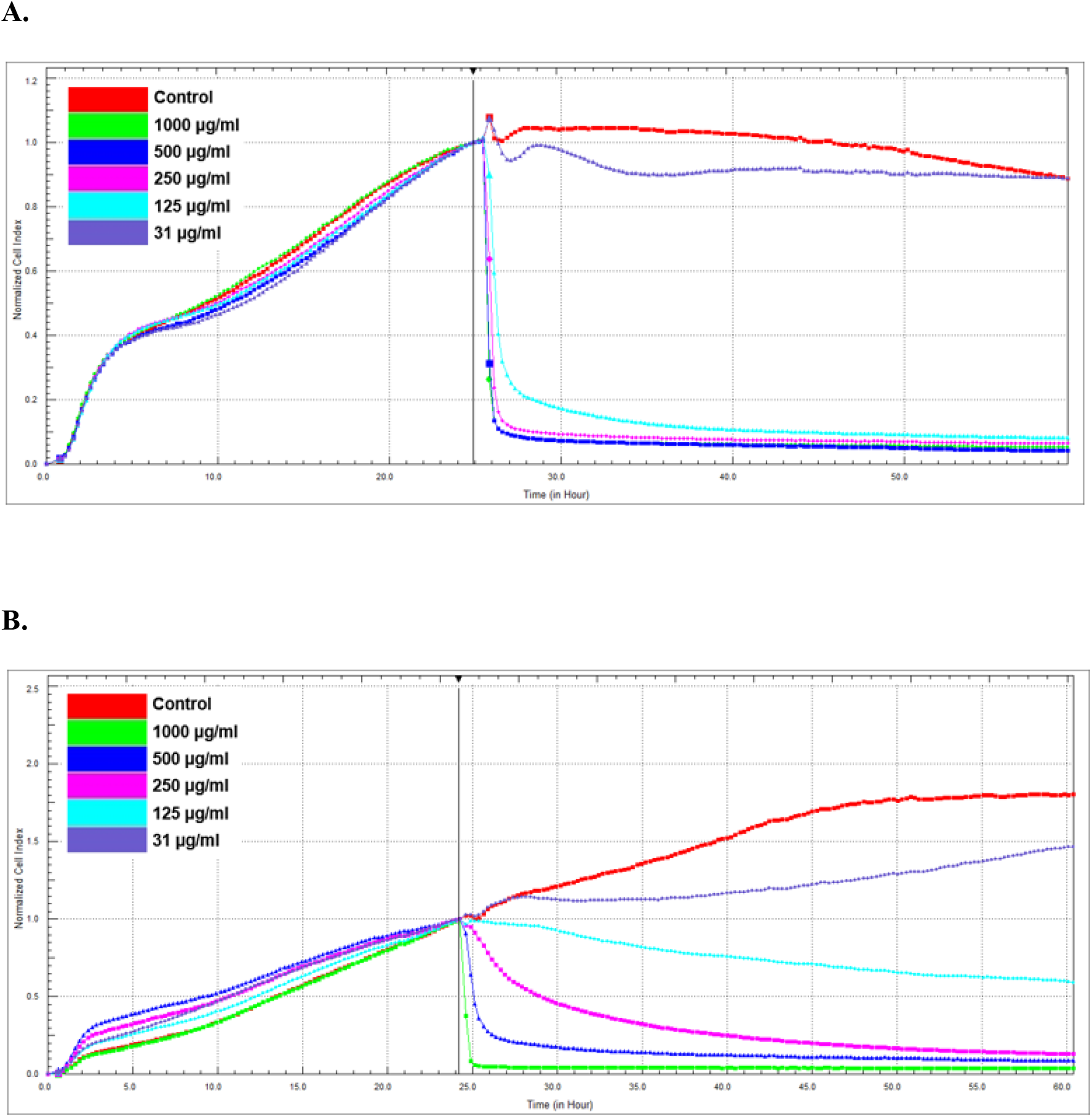
Proliferation rate of UT-SCC-74A cells incubated with different concentrations of either KK **(A)** or KS **(B)** propolis extracts. The rate of proliferation was monitored in realtime until the growth of the control reached the plateau. RTCA Software version 1.2 was used for data analysis. IC_50_ of KK was calculated as 39,12 μg/ml and IC_50_ of Sakyatan sample was calculated as 119,36 μg/ml at 24 hour following the propolis administration.

As a next step, we investigated the effect of two propolis samples on migration capacity of UT-SCC-74A primary head and neck cancer cell line. For this purpose, we utilized xCELLigence real time cell analyzer system with CIM plates. CIM-plates contain two chambers separated by a microporous membrane able to detect the cells migrated by the microelectrodes attached to the bottom part of the membrane.10% fetal bovine serum (FBS) was used as chemoattractant and serum-free medium was used as negative control. Both KK and KS propolis effectively inhibited the migration of cancer cells at their previously determined IC_50_ dosages (Figure 2).

**Figure 2.**
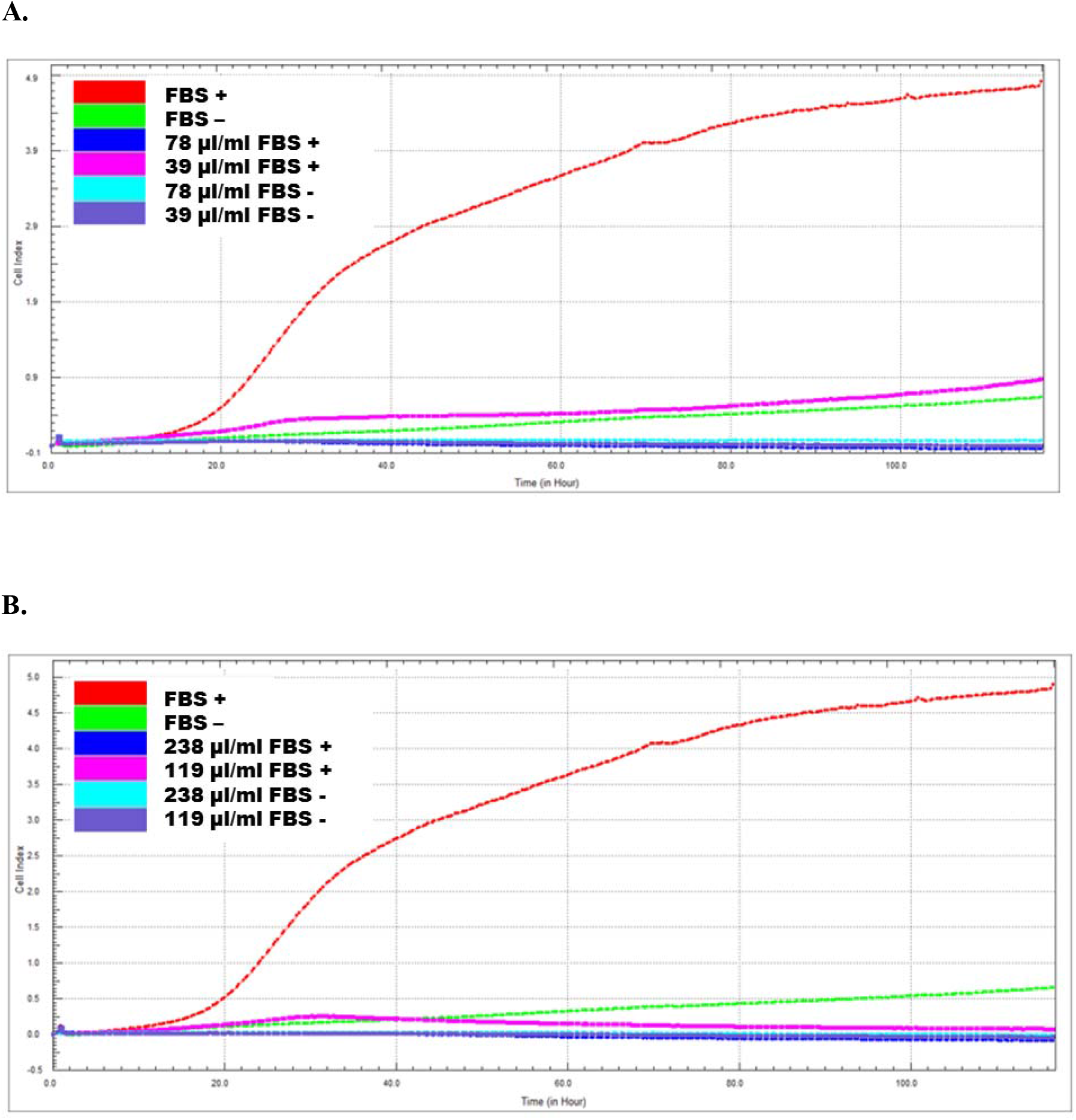
Migration of UT-SCC-74A cells after treatment with IC50 concentrations of KK **(A)** and KS **(B)** propolis. Migration was monitored in real-time by using xCELLigence technology.

Different mechanisms have been indicated as underlying reason for anticancer properties of propolis. Several studies showed that propolis can trigger both intrinsic and extrinsic apoptotic pathways in cancer cells and this effect is depend on the cell type (Wang et al. 2010; Orsolic et al. 2004; Seda Vatansever et al. 2010; Motomura et al. 2008; Eom et al. 2010). Another main effect of propolis is suppressing tumor cell proliferation by either inducing cell cycle arrest or decreasing telomerase activity (Motomura et al. 2008; Gunduz et al. 2005).

Even though both KK and KS propolis can effectively inhibit tumor cell proliferation and migration at different concentrates, further studies needed to evaluate the molecular mechanism behind the antitumor effect observed.

## 3. Experimental

### 3.1. Propolis material and preparation of the extracts

Propolis materials were collected from Konya Sakyatan (KS) and Kiziloren (KK) regions. The samples (5 g) were macerated with methanol (100 mL) at room temperature for 24 h. Extracts were filtered through a filter paper. Solvent from the samples was removed using a rotary evaporator.

### 3.2. Quantification of phenolics content by RP-HPLC

Phenolic compounds were evaluated by RP-HPLC (Shimadzu Scientific Instruments, Tokyo, Japan). Detection and quantification were carried out with LC-10ADvp pump, a Diode Array Detector, a CT0-10Avp column heater, SCL-10Avp system controller, DGU-14A degasser and SIL-10ADvp auto sampler (Shimadzu Scientific Instruments, Columbia, MD). Separations were carried out at 30°C on Agilent^®^ Eclipse XDB C-18 reversed-phase column (250 mm × 4.6 mm length, 5 μm particle size). The eluates were detected at 280 nm.

Phenolic compositions of the extracts were determined by a modified method of Sarikurkcu et al (2014). Gallic acid, (+)-catechin, p-hydroxybenzoic acid, chlorogenic acid, caffeic acid, (−)-epicatechin, ferulic acid, benzoic acid, rutin, rosmarinic acid, and apigenin were used as standards. The phenolic compounds were characterized according to the UV spectra, retention times, and comparison with authentic standards. The identified phenolic acids were quantified by comparison of the area of their peaks recorded at 280 nm with calibration curves obtained from commercial standards of each compound. The results were expressed in mg per g of dry extract.

### 3.3. Total bioactive compounds

The total phenolic content was determined as previously reported by Zengin et al., (2014), the results being expressed as milligrams of gallic acid equivalents (mg GAE/g extract). The total flavonoid content was determined using the method described previously by Ceylan et al., (2016) and Lazarova et al., (2014). Rutin was used as reference standard and the results were expressed as milligrams of rutin equivalents (mg RE/g extract).

### 3.4. Antioxidant activity

The DPPH radical scavenging and ABTS radical cation scavenging activities were determined using the previous described method (2014), and results were expressed as trolox equivalents (TEs/g extract).

The reducing power was measured according to the method reported by Zengin et al. (2014), using cupric ion reducing antioxidant power (CUPRAC) and ferric ion reducing antioxidant power, and results were expressed as trolox equivalents (TEs/g extract).

Total antioxidant capacity was determined using phosphomolybdenum and β-carotene/linoleic acid bleaching methods (2014). Metal chelating activity on ferrous ions, determined by the method described by Zengin et al (2014) was expressed as EDTA equivalents (EDTAEs/g extract).

### 3.5. Cell Culture

UT-SCC-74A (University of Turku Squamous Cell Carcinoma) cells were a kind gift from Prof. Reidar Grenman (Department of Otorhinolaryngology, University of Turku and Turku University Hospital, Turku, Finland). These cells were generated from head and neck primary tumors. UT-SCC-74A cells were cultured in Dulbecco’s modified Eagle’s medium (DMEM Hyclone, Logan, USA) containing 10% fetal bovine serum (FBS, Hyclone), L-glutamine (Hyclone) and penicillin/ streptomycin (Hyclone) at 37°C in a humidified atmosphere of 5% CO2 in air.

### 3.6. Proliferation Assay with the xCELLigence System

Cell proliferation assay with using the xCELLigence real time cell analyzer system (ACEA Bioscience, San Diego, USA) as we previously described (Lazarova et al. 2015). Briefly 100 μL of medium was added to each well and a one minute calibration measurement was taken. Right after, 1 x 10^4^ UT-SCC-74A cells in 100 μL of medium were added to each well. The system was then set to take measurements every 15 minutes for 24 hours. After 24 hours, the medium was removed and replaced with medium containing either KK Propolis or KS propolis methanolic extracts with 0,1% DMSO (1000, 500, 250, 125, 31 μg and DMSO-only control). Each dose was performed in triplicate. The system was then set to take measurements every 15 minutes for 36 hours. Data analysis was performed using RTCA Software version 1.2. IC_50_ values (% mL-1) were calculated based on the final readings taken at the 60 hour time point. Six concentrations were used in cell treatment and IC50 calculations.

### 3.7. Migration Assay with the xCELLigence System

xCELLigence real time cell analyzer system (ACEA Bioscience, San Diego, USA) was used to investigate cell migration in UT-SCC-74A cells in a label-free environment. Migration was examined on 16-transwell CIM plates with microelectrodes attached to the underside bottom of the membrane for impedance-based detection of the migrated cells.

First, 160 μL of medium either containing 10% FBS (as chemoattractant, FBS +) or serum-free (as negative control, FBS-) was added to all wells of the lower chamber. CIM-16 plates were further prepared according to the manufacture’s protocol. Background signals generated by the cell-free media were recorded. Cells were harvested using trypsin, counted and resuspended in 238 and 119 ug/ml propolis an appropriate volume of DMEM containing either 10% FBS or only DMEM. Cells (100 000 cells per 37 μl medium) were seeded onto the upper chamber of the CIM-16 plate and allowed to settle onto the membrane. The programmed signal detection for quantification of the cell index was measured every 15 min over a period of 120 h.

## 4. Conclusion

Propolis components can greatly vary from one sample to another even in the same region, hence propolis selections for cancer prevention and treatment studies should be carefully considered.

